# A study on tomato brown rugose fruit virus longevity in soil and virion susceptibility to various pH treatments contributes to optimization of a soil disinfection protocol

**DOI:** 10.1101/2024.01.29.577727

**Authors:** Ori Molad, Elisheva Smith, Neta Luria, Elena Bakelman, Oded Lachman, Meital Reches, Aviv Dombrovsky

## Abstract

**Background and aims:** Tobamoviruses are highly stable soil-borne pathogens posing a challenge to a monoculture practice. Biochemical and physical properties of tobamovirus virions were studied by analyses of tobacco mosaic virus (TMV). Little is known about tomato brown rugose fruit tobamovirus (ToBRFV) regarding longevity in soil and virion stability. Our aims were to determine ToBRFV longevity in naturally-contaminated soil and study virion stability in a range of acidic and alkaline conditions to promote new strategies for soil remediation.

**Methods:** ToBRFV longevity in naturally-contaminated soil was tested by collecting an earth pile after a growth-cycle of ToBRFV-infected tomato plants. The soil was sampled at different time points and root-truncated tomato seedlings were planted. Virion stability at a range of pH values was determined by testing virus infectivity on *Nicotiana glutinosa*; by amplifying large genome segments using RT-PCR; and by transmission electron microscopy (TEM) visualization.

**Results:** ToBRFV-infectivity in naturally-contaminated soil was profoundly reduced by day 184 of pile-age and was abolished between 205-385 days of pile-age. Virion stability and genome integrity were preserved over the pH range of 2-10. At pH 1, ToBRFV-infectivity and efficiency of large genome segment amplifications were reduced. At pH values above 10, modified particle morphologies were visualized by TEM, and virus infectivity was abolished. Treatment of ToBRFV-contaminated soil with an alkaline chlorinated-trisodium phosphate solution profoundly reduced soil-mediated virus infection of root-truncated tomato seedlings.

**Conclusions:** pH values above 10, compromised ToBRFV particle morphology genome integrity and virus infectivity. Alkaline disinfectant enhanced soil remediation following natural ToBRFV contamination.

## Introduction

Tomato brown rugose fruit virus (ToBRFV) is a positive sense RNA virus belonging to the genus *Tobamovirus*, family *Virgaviridae*. The ToBRFV virion is a rod-shaped particle of approximately 18 nm in diameter and 235±123 nm in length (Luria et al., 2017). The single stranded positive sense RNA genome is about 6,400 nucleotides long, encoding the four ORFs of two replicase proteins, the coat protein (CP) and the movement protein (MP) (Luria et al., 2017). Since it was first identified in 2014 in commercial tomato (*Solanum lycopersicum* L.) greenhouses in Israel and Jordan (Luria et al., 2017; Salem et al., 2016), the virus spread to other countries in the Middle East, Europe, and Central and North America, hindering tomato production (Zhang et al., 2022). For decades, the *Tm-2^2^* resistance allele provided commercial tomato cultivars broad resistance against tobamoviruses (Lanfermeijer et al., 2004; Pelham, 1966). However, ToBRFV overcomes *Tm-2^2^* resistance (Luria et al., 2017). Mutations in MP of tobacco mosaic virus (TMV) and tomato mosaic virus were shown to overcome *Tm-2* and *Tm-2^2^* resistance alleles, respectively and similarly the resistance breaking of ToBRFV has been recently attributed to the viral MP (Hak & Spiegelman, 2021; Meshi et al., 1989; Weber et al., 1993; Yan et al., 2021).

ToBRFV worldwide spread and distribution has prioritized tomato industry research on virus epidemiology, seed transmission, diagnostics and breeding strategies in search of resistant varieties (Zhang et al., 2022; Zinger et al., 2021). Tobamoviruses are highly stable and persist under extreme environmental conditions. Tobamoviruses are very well preserved in plant debris, soil and agricultural tools such as trellising ropes and surfaces in the greenhouse infrastructure and workers’ facilities (Broadbent & Fletcher, 1963; Broadbent et al., 1965; Lanter et al., 1982). TMV remained infective in buried root debris for at least 13 months (Broadbent et al., 1965; Broadbent, 1976). The cucurbit-infecting tobamovirus cucumber green mottle mosaic virus (CGMMV) remained infective in soil for 17 months after the removal of infected watermelon plants (Lovelock et al., 2022). In addition, overwintered soil contaminated with CGMMV-infected plant debris remained infective in bottle gourd propagation (Li et al., 2016). The contaminated soil serves as a primary source of infection occurring *via* root-damaged seedlings, which are predisposed to soil-mediated viral infection (Broadbent, 1965; Dombrovsky et al., 2022).

Tobamovirus persistence in soil depends on various factors including clay content, organic matter, ionic strength and pH (Williamson et al., 2017). Ionic strength and soil pH are negatively correlated with high virus abundance in soil by affecting virus attachment to various surfaces (Gerba, 1984; Loveland et al., 1996; Williamson et al., 2017; Yoshimoto et al., 2012). Several soil disinfection strategies such as solarization and steaming, as well as the application of commercially available disinfectants to inactivate tobamoviruses were efficient in reducing virus infectivity (Chanda et al., 2021; Darzi et al., 2020; Dombrovsky et al., 2022; Ling et al., 2022; Luvisi et al., 2015; Panth et al., 2020; Smith & Dombrovsky, 2019; Vargas-Mejía et al., 2023). However, mechanisms of disinfectant effects on ToBRFV stability have not been studied yet. In addition, we have recently shown that soil-mediated transmission of ToBRFV occurred under natural contamination conditions, following a growth cycle of virus-infected tomato plants, and the virus infectious potential was apparently dependent on the length of the growth cycle of the infected crop (Klein et al., 2023). However, ToBRFV longevity in soil under the natural contamination conditions has yet not been established.

The current study aimed to investigate perseverance of ToBRFV infectious potential in naturally contaminated soil and to characterize virion stability under acidic and alkaline conditions with respect to particle morphology, genome integrity and infectivity. Such data could be used to establish improved soil disinfection protocols.

## Materials and Methods

### ToBRFV infectivity in soil collected after a growth cycle of virus-infected tomato

Soil of naturally infected tomato plants was collected from two commercial sites: Ramat Negev in southern Israel and Ahituv in the Sharon plain in central Israel. The soil was subjected to virion purification and the virome visualized by transmission electron microscopy (TEM) (see below). ToBRFV-infected plants were grown in a commercial soil medium Green 90 (EvenAri, Beit Elazari, Israel) for 6 months to resemble a growth cycle. At the end of the growth cycle, the soil was cleared of plant debris excluding the roots; then the soil was collected and piled up. The soil was tested for the presence of ToBRFV by virion purification, large segment (LS) RT-PCR amplification and western blot analysis using specific antibodies against ToBRFV (see below) (Luria et al., 2017). Two soil piles were accumulated in a net house: one was kept dry and the second was irrigated twice daily for 20 min and mixed every 15 days, allowing weed cover to grow. Infectivity potential tests of ToBRFV from the dry and wet soil piles were conducted for 79 and 385 days, respectively. On each testing day, samples from the piles were dispersed into pots and the infectious potential was tested by planting 49 to 57 root-truncated tomato plants cv. Ikram harboring the *Tm-2^2^* resistance allele. Plants were analyzed for ToBRFV infection 30 days post-planting using enzyme linked immunosorbent assay (ELISA) (see below).

### ToBRFV virion purifications from soil

ToBRFV virion purification from soil samples was conducted by suspending soil in sodium phosphate buffer (0.1M, pH=7.0). The suspension was stirred at room temperature for 24 h, and then filtered through Miracloth (Merck, USA). The suspension was then centrifuged for 20 min at 9,700 *g* at 6°C using an Avanti J-E centrifuge and a JLA-16.250 rotor (Beckman Coulter, USA). The supernatant was separated and centrifuged for 3 h at 200,000 *g* at 6°C using an Optima XPN-90 ultracentrifuge with a Type 45 Ti rotor (Beckman Coulter, USA). The supernatant was discarded, and the pellets were suspended overnight at room temperature with 500 µl sodium phosphate buffer (0.01M, pH=7.0). For further purification and increasing virion concentration, aliquots of 400 µl of the suspended pellet were added to 1.2 ml 20% sucrose in sodium phosphate buffer (0.01M, pH=7.0). The two-phased solutions were centrifuged for 1 h at 10,000 *g* at 4°C. The supernatant was transferred to 5.2 ml vials with the addition of an equal volume of sodium phosphate buffer (0.01M, pH=7.0), to reduce sucrose concentrations to ∼10%, and centrifuged for 3 h at 260,000 *g* using an SW 55 Ti swinging-bucket rotor (Beckman Coulter, USA). The final supernatant was discarded, and the obtained pellets were suspended overnight at 4°C with 100 µl sodium phosphate buffer (0.01M, pH= 7.0).

### Virion purifications from ToBRFV-infected tomato plants

Tomato plants cv. Ikram were dusted with carborundum and hand-rubbed with a ToBRFV inoculum solution, prepared from infected plant debris homogenized in water. After the emergence of foliar symptoms, 200 g of symptomatic tomato leaves were collected and homogenized in 400 ml sodium phosphate buffer (0.1M, pH=7.0). Forty ml each of chloroform (stab. Amylene) and *n*-butanol were added, and the mixture was stirred for 1 h at 4°C. The mixture was then centrifuged for 20 min at 9,700 *g* at 6°C using an Avanti J-E centrifuge and a JLA-16.250 rotor (Beckman Coulter, USA). The supernatant was separated and centrifuged for 2.5 h at 200,000 *g* at 6°C using an Optima XPN-90 ultracentrifuge and a Type 45 Ti rotor (Beckman Coulter, USA). The supernatant was discarded, and the pellets were suspended overnight at 4°C with 1 ml sodium phosphate buffer (0.01M, pH=7.0).

### pH-modified ToBRFV virions and virion RNA solutions

To prepare pH-modified stock solutions, NaOH (1M) or HCl (1M) was added to 0.01M sodium phosphate buffer with a starting ionic strength of 0.04M (Fig. S1). To prepare pH-modified virions or virion RNA (see below), 1 µg/µl ToBRFV purified virions or 2 µg/µl virion RNA were diluted 1:10 with each stock solution and incubated for 1 h at 23-25°C. The final pH of each solution was measured, and no pH deviations were observed. In addition, ToBRFV virions were treated with chlorinated-trisodium phosphate (Cl-TSP) solutions of 1% (pH=11.43), 3% (pH=11.6), 5% (pH=11.6), and 3% Cl-TSP + 1% KOH (pH=12.47) as well as NaOH (pH=13), solutions that were used for soil disinfection studies as well (see below).

### Virion RNA extraction and reverse transcriptase polymerase chain reaction (RT-PCR)

Virion RNA was extracted from ToBRFV virions using the AccuPrep Viral RNA Extraction Kit (Bioneer, Republic of Korea), according to the manufacturer’s instructions. For RT-PCR, viral RNA served as a template for cDNA synthesis using Maxima First Strand cDNA Synthesis Kit for RT-qPCR (Thermo Fisher Scientific, USA) with the reverse primer R-6,392: 5’-TGGGCCCCTACCGGGGGTTCCG-3’ designed for ToBRFV 3’ UTR. The cDNA obtained served as a template for PCR amplifications using Platinum™ SuperFi II Green PCR Master Mix (Thermo Fisher Scientific, USA) and specific primers for ToBRFV LS PCR amplifications: F-1: 5’-GTGTATTTTTTACAACATATACCAAC-3’ and R-6,392 for ∼6.3 kb amplicons and F-264: 5’-AGGGCATATCCAGAATTCCA-3’ and R-6,332: 5’-ATGTGTATGAACCATACACATTTGTC -3’ for ∼6 kb amplicons. In addition, a ToBRFV specific primer pair (F-5196 5’-GGAGAGAGCGGACGAGGCAA-3’ and R-5878 5’-ACAGGTTTCCACACTTCGCT-3’) was used to obtain a 682 bp segment. Amplicons were Sanger sequenced at the ToBRFV CP region.

### Biological assays of pH-modified ToBRFV virion and virion RNA solutions

*Nicotiana glutinosa* L. plants were inoculated with pH-modified virions (∼7 µg/leaf) and virion RNA solutions (∼5 µg/leaf) to assess quantitatively the effect of pH on ToBRFV infectivity (Luria et al., 2017). Two leaves per plant (1 plant per treatment) were inoculated with each pH-treated virion solution and one leaf per plant (3 plants per treatment) was inoculated with each pH-treated virion RNA solution.

### Indirect enzyme-linked immunosorbent assay (ELISA)

Tomato leaves were collected, ground in coating buffer (∼0.5 g/1 ml), and tested using indirect ELISA (Clark & Adams, 1977). Samples were analyzed in duplicates. Laboratory-produced ToBRFV antiserum (Luria et al., 2017), diluted to a ratio of 1:4,000 in PBS-milk (2% non-fat milk powder in PBS), was added to the samples. Plates were incubated for 2 h at 37°C or overnight at 4°C. Commercial alkaline phosphatase (AP)-conjugated goat anti-rabbit (IgG) antibodies (Sigma-Aldrich, USA, diluted 1:5,000 in PBS) were added and incubated for 2 h at 37°C. p-nitro phenyl phosphate substrate (Sigma) was used at a concentration of 0.6 mg/ml for AP activity detection measured at 405 nm and 620 nm. Samples with optical density (OD) values of 2.5-3 times the negative control were considered ToBRFV positive.

### Negative staining and transmission electron microscopy (TEM) visualization of ToBRFV

For TEM analysis, 2.5 µL of pH-modified ToBRFV virions, Cl-TSP-treated ToBRFV virions or soil virion preparations were applied to Formvar/Carbon 300 mesh copper support grids (Ted Pella Inc., USA) for 30 sec. After blotting the excess liquid, a drop of 2.5 µl of 2% (w/v) ammonium molybdate, 0.1% (w/v) trehalose solution (pH=7.2) was applied for 30 sec for negative staining. The remaining excess fluid was blotted, and the grids were allowed to dry for 1 h at room temperature. The samples were visualized using a Tecnai G2 Spirit Twin TEM at 120 kV accelerating voltage (FEI, USA).

### Soil disinfection tests of ToBRFV-contaminated soil

To obtain ToBRFV-contaminated soil, ToBRFV-inoculated tomato seedlings were grown in a net house for 4-6 months. Plant infections were confirmed using ELISA. At the end of the growth cycle, plants were removed, and the soil was collected, mixed, and distributed into five 10 L pots. To test the effect of alkaline pH on ToBRFV-infected soil, four different treatment solutions were prepared, two based on Cl-TSP (97% trisodium phosphate, 3% chlorine). Crystalline Cl-TSP is a complex structure of sodium hypochlorite (NaOCl), trisodium phosphate (TSP; Na_3_PO_4_) and water. The Cl-TSP complex is stable compared to NaOCl. In aqueous solutions, OCl^-^ ions are in equilibrium with hypochloric acid (HOCl), which is unstable. High pH conditions (provided either by NaOH or Na_3_PO_4_, or both) shift the equilibrium toward OCl^-^ and thus inhibit hypochlorite degradation. The four soil treatments were: tap water (pH=8.35), 3% Cl-TSP (pH=11.6), 3% Cl-TSP + 1% KOH (pH=12.47), and tap water with added NaOH (pH=13.0). Each treatment was prepared in 5 L and was poured gradually into the 10 L ToBRFV-contaminated soil buckets while the fifth pot served as an untreated positive control. The tap water treatment served as a control for water-mediated virion displacement. The four solutions were incubated with the ToBRFV-contaminated soil for 24 h. Following the treatments, soil was sampled for western blot analyses (see below) and the soil buckets were washed three times with 5 L tap water to alleviate the effects of alkaline treatments on soil porosity and structure before testing the soil for ToBRFV infectivity. Remediated soils and the untreated soil were then distributed into 0.3 L pots. Root-truncated tomato seedlings were planted in the pots, 20-25 plants per each treatment, and at 21 days post planting ToBRFV detection was conducted using ELISA. Three experiments were conducted. For statistical analyses the Two Sample t-Test assuming unequal variances was used.

### SDS-PAGE, Coomassie blue staining and western blot analyses

Soil virion preparations were subjected to total protein extractions using the urea-sodium dodecyl sulfate (SDS)-ß-mercaptoethanol extraction buffer (75 mM Tris–HCL pH 6.8, 9 M urea, 4.5% SDS, 7.5% ß-mercaptoethanol). The products of the extractions were added to Laemmli loading buffer (Laemmli, 1970). In addition, equally sampled pH-modified ToBRFV virions or ToBRFV virions subjected to treatments with the various Cl-TSP solutions were added to Laemmli loading buffer. The protein samples were separated on 15% SDS– polyacrylamide gel electrophoresis (PAGE) for 30 min at 40 V followed by 1.5 h at 100 V. Gels were either subjected to Coomassie blue staining or a western blot analysis. For the western blot, gels were electro-blotted onto a nitrocellulose membrane for 30 min at 200 mA using a semidry transfer apparatus (Bio-Rad, California, USA). Membranes were blocked with 3% non-fat dry milk in PBS for 2 h and then incubated with ToBRFV-specific laboratory-produced primary antibodies (1:4000) for 40 min at room temperature and overnight at 4 °C. After four washes with PBS-T (0.05% v/v Tween-20 in PBS), the membranes were incubated with goat anti-rabbit AP-conjugated antibodies (1:5000 in PBS, Sigma-Aldrich, Missouri, USA), followed by four washes with PBS-T. An AP substrate (NBT/BCIP, Bio-Rad, California, USA) was used to detect ToBRFV CP. In the case of Cl-TSP treated soil, 24 h after soil treatments with Cl-TSP and before soil remediation, 30 g of the treated soils were equally sampled. Ten ml of double distilled water were added to the dry, untreated soil from the control group. The soil mixtures were centrifuged for 15 minutes at 6,000 rpm at 6°C using a Hermle Z 326 K centrifuge and a 221.55 V20 adapter for 50 ml vials (Hermle Labortechnik, Germany). The supernatant was removed, centrifuged again, and added to Laemmli loading buffer and run on SDS PAGE followed by a western blot.

## Results

### ToBRFV was preserved in soil virome at the end of a growth cycle of ToBRFV-infected tomato plants

Soil samples were collected at the end of a tomato growth cycle in two commercial greenhouses in Israel: Ramat Negev in southern Israel and Ahituv in the Sharon plain of central Israel. The tomato plants were naturally infected with ToBRFV and all aerial plant material was removed (Fig. 1a). Tobamovirus rod-like particles were apparent in analyses of virion purifications from the soil samples when the soil virome was visualized by TEM, (Fig. 1b-d). Soil ToBRFV was identified by amplifying large genome segments of ∼6.3 kb and ∼6 kb that were Sanger sequenced at the ToBRFV CP region (Fig. 1e, f). In addition, a western blot analysis using specific antibodies raised against ToBRFV CP (Luria et al., 2017) showed the presence of ToBRFV in the soil (Fig. 1g).

**Fig. 1.**
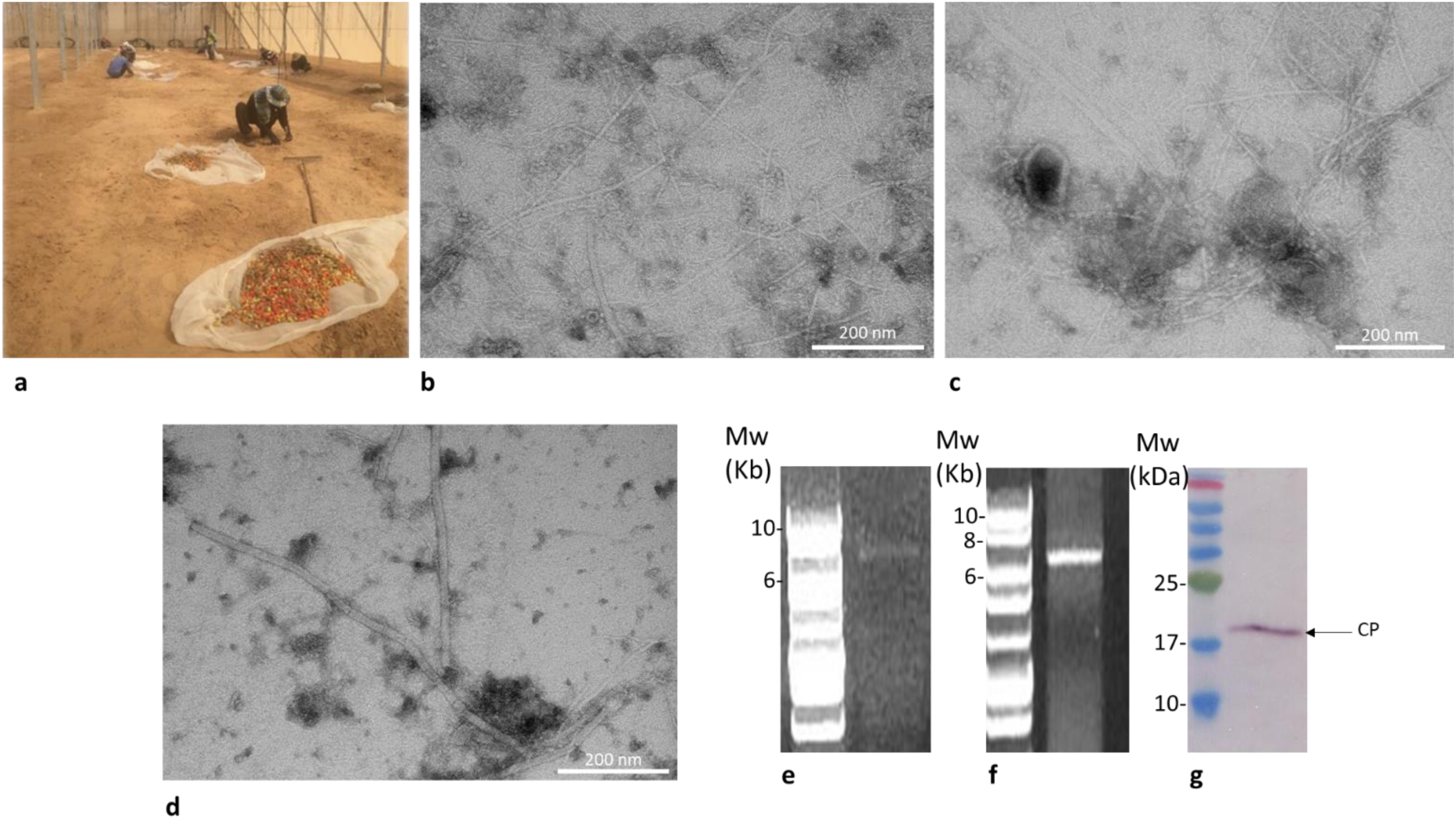
ToBRFV contaminated soil in commercial greenhouses after a growth cycle of naturally infected crops (a) Greenhouse workers in Ramat Negev, Israel, remove fruit and plant debris after a growth cycle. (b, c) Representative TEM micrographs of virions purified from soil of a naturally infected crop collected from Achituv, Israel. (d) A representative TEM micrograph of virions purified from soil of a naturally infected crop collected from Ramat Negev, Israel. (e, f) LS RT-PCR amplification of ∼6.3 kb and ∼6 kb large fragments of ToBRFV in soil virions using the primer sets F-1 and R-6,392 and F-264 and R-6,332, respectively that were Sanger sequenced at the ToBRFV CP region. (g) A western blot analysis of soil virion purification detected ToBRFV CP of ∼17.5 kDa using specific antiserum.

### ToBRFV longevity in naturally contaminated soil

We have recently demonstrated that ToBRFV in naturally contaminated soil was infectious and infectivity potential was dependent on length of the growth cycle of the infected crop (Klein et al., 2023). For studying ToBRFV longevity in naturally contaminated soil, we piled up soil from a six-month growth cycle of ToBRFV-inoculated tomato plants to achieve a significant soil-inoculum equivalent to the contaminated soil of a commercial growth cycle (Fig. 2a). In our model, two different piles were studied: a dry pile, resembling fallowed soil and a wet pile, irrigated twice daily to enhance microbial activity and decomposition of organic matter (Fig. 2b, c). Soil was sampled from the piles at various days of pile-age to test ToBRFV infectivity. For the infectious potential tests, the study was conducted under stringent conditions by performing root truncation prior to seedling planting in order to increase root susceptibility to ToBRFV soil-mediated infection (Fig. 2d). Leaf samples from the planted tomato seedlings of each time-point of pile-age were collected at 40 days post planting and analyzed using ELISA. ToBRFV positive plants were symptomatic (Fig. 2e). ToBRFV longevity in the dry and wet earth piles was tested for 79 and 385 days, respectively. ToBRFV in the dry earth pile showed a peak of 20% soil-mediated infectivity at the 9^th^ day of pile accumulation, and by the 79^th^ day soil infectivity was reduced to 2% (Fig. 2f). However, ToBRFV in the wet earth pile showed higher infectivity of 40-50% between 54-98 days of pile-age, and reduction to 3% infectivity was observed by day 184 of pile-age (Fig. 2g). These differences could reflect the decomposition of organic matter enhanced in the wet pile that apparently has also assured zero percent infectivity observed between 205-385 days of pile-age.

**Fig. 2.**
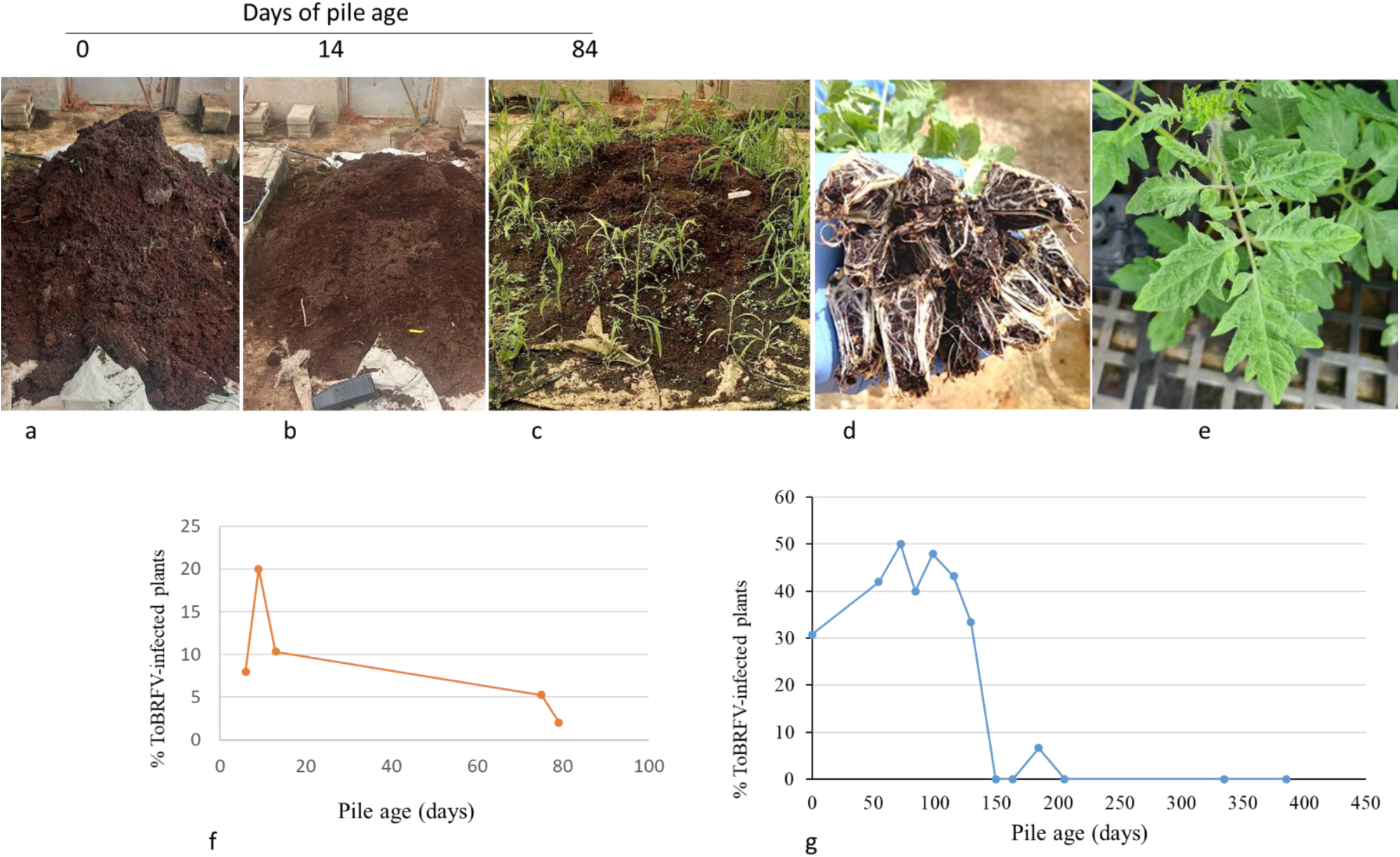
ToBRFV longevity in naturally contaminated soil collected after a 6-month growth cycle of virus-infected tomato plants (a) A pile of naturally contaminated soil collected after a 6-month growth cycle of virus-infected tomato plants. (b) Sampling at 14 days of wet-pile age. (c) A pile kept wet, allowing growth of a cover crop. (d) Root truncation of tomato seedlings before planting to test ToBRFV soil-mediated infectivity. (e) Symptoms of severe mosaic in ELISA positive plants. (f) ToBRFV-infectious potential during 79 days of the dry earth pile. (g) ToBRFV infectious potential during 385 days of the wet earth pile.

### Extreme acidic and alkaline conditions modified ToBRFV virion morphology as visualized by TEM

To enhance ToBRFV inactivation in soil we studied ToBRFV virion stability under a range of acidic and alkaline conditions. Purified ToBRFV virions were incubated at different pH conditions for an hour at room temperature (23-25°C) and were then visualized by TEM (Fig. 3). Rod-like structures characteristic of tobamovirus particle morphology were observed at pH values of 1-10, with a low abundance observed of particles at pH 1 (Fig. 3a-j). At pH 11 truncated rod-shaped virion structures were apparent of up to ∼200 nm in length with abundant discs (Fig. 3k). No distinct structures were observed at pH 12 (Fig. 3l).

**Fig. 3.**
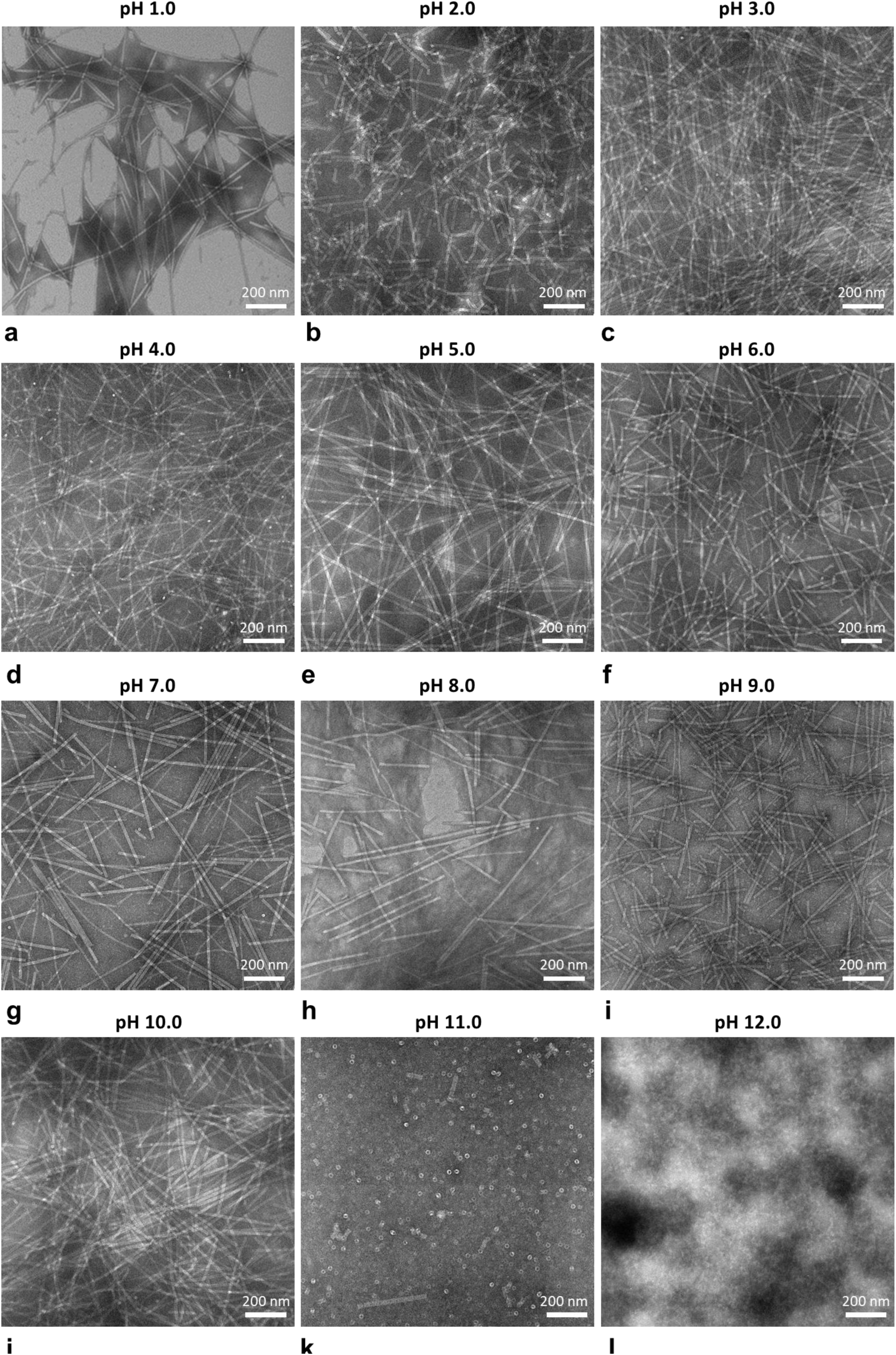
Representative TEM micrographs of pH-modified ToBRFV virions. (a-j) Rod like particles characteristic of tobamoviruses were visualized in pH-modified virions over the pH range 1-10. (k) Truncated virions of up to ∼200 nm and structures resembling 20S discs were visualized in virion solutions incubated at pH 11. (l) Undefined structures were observed in virion solutions incubated at pH 12.

### Infectious potential of ToBRFV virions and virion RNA was reduced under extreme acidic and alkaline conditions

The infectious potential of the pH-treated ToBRFV virions was examined using a biological assay of local lesion (LL) expression on *N. glutinosa* plants. At pH 1, the average LL counts per leaf was significantly lower than the average LL counts per leaf observed at pH values of 2-10. At pH values of 11 and 12 no LL were observed (Fig. 4a, b). In addition, virion RNA was treated with the various pH solutions and the pH-modified virion RNA was tested for infectivity on *N. glutinosa* plants. pH-treated virion RNA showed no infectivity potential at pH 1 and LL counts were already reduced at pH 10 (Fig. 4a).

**Fig. 4.**
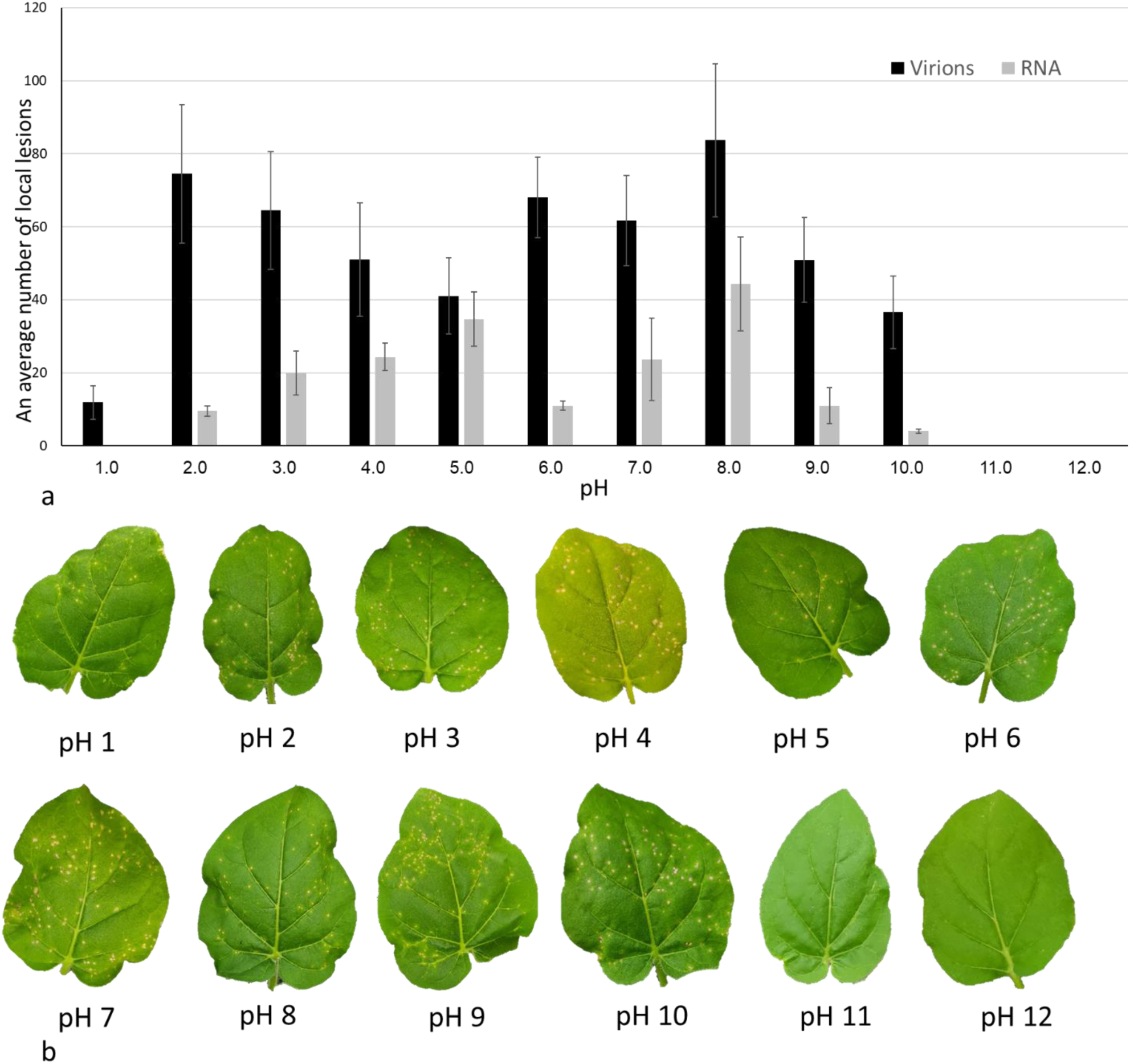
Extreme acidic and alkaline solutions reduced the infectious potential of ToBRFV virions and virion RNA. (a) Local lesion (LL) counts of pH treated virions and virion RNA using *N. glutinosa* plants. Bars represent SEM. (b) Sampled *N. glutinosa* leaves showing local lesions of pH-modified virions.

### Genome integrity was compromised at extreme acidic and alkaline conditions

ToBRFV virions and RNA extracted from ToBRFV virions subjected to pH treatments were analyzed for genome integrity by LS RT-PCR of the ∼6.3 kb genome segment (Fig. 5a1, b1). RT-PCR of a ∼0.7kb genome segment was performed as a control for the presence of ToBRFV RNA (Fig. 5a2, b2). LS amplification was obtained at pH 3-10 in samples of both the pH-modified ToBRFV virions (in 4-7 reactions of 8) and pH-modified ToBRFV virion RNA (in 3-4 reactions of 4) (Table 1). At pH values above and below that range, LS amplifications were not readily obtained using pH-modified virion samples (in 1-3 reactions of 8) (Table 1). ToBRFV virion RNA showed higher sensitivity to treatments with the extreme acidic and alkaline solutions showing no large genome-fragment amplifications (in four different reactions) at pH values of 1, 2 and 12 (Fig. 5b1, Table 1).

**Fig. 5.**
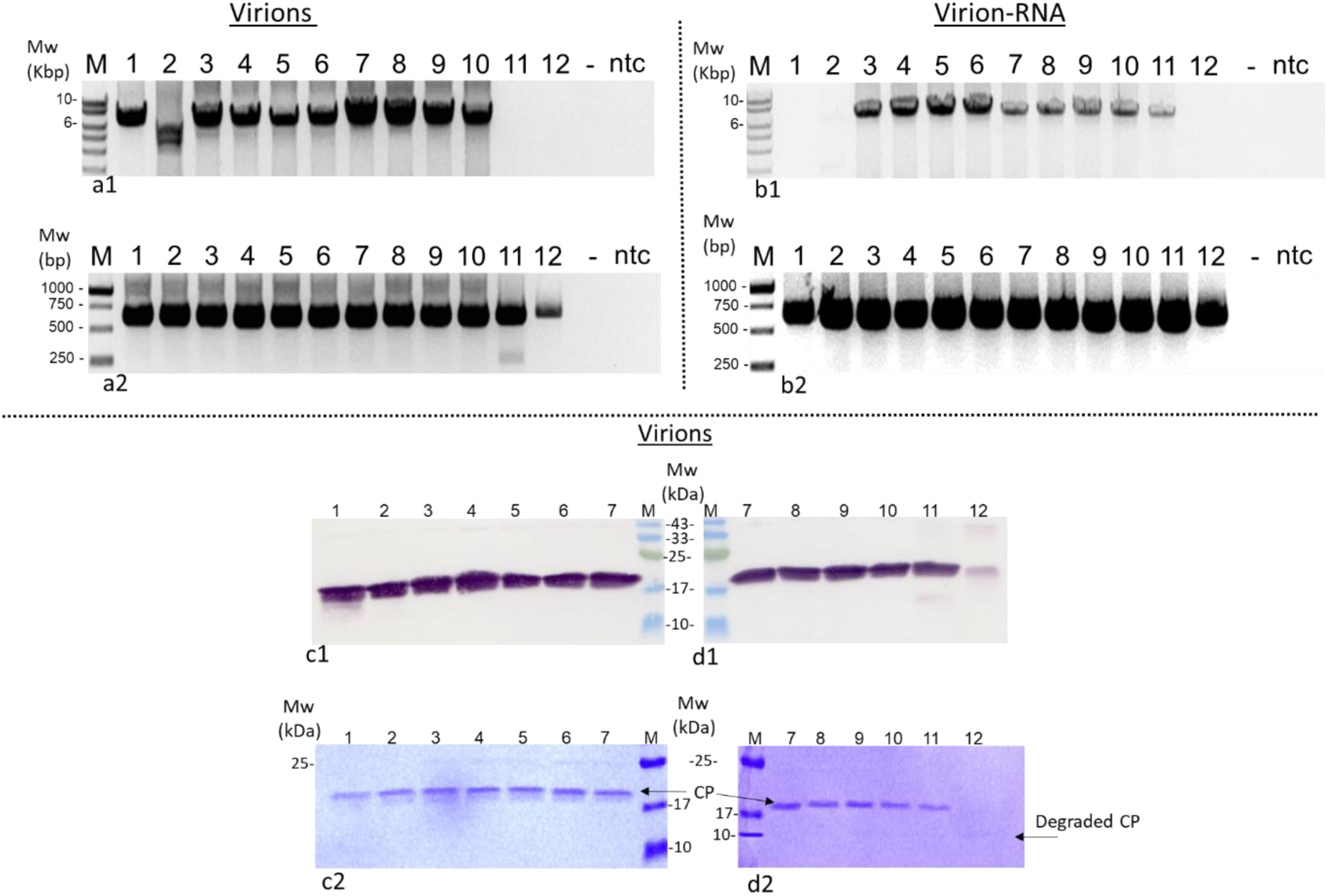
ToBRFV compromised genome integrity and CP degradation occurred under extreme alkaline conditions. (a1) pH-modified virions subjected to LS RT-PCR amplifications of a large ∼6.3 kb fragment using the primer pair F-1 and R-6,392. (a2) pH-modified virions subjected to RT-PCR amplifications of a small ∼0.7 kb fragment using the primer pair F-5,196 and R-5,878. (b1) pH-modified virion RNA subjected to LS RT-PCR amplifications of a large ∼6.3 kb fragment. (b2) pH-modified virion RNA subjected to RT-PCR amplifications of a small ∼0.7 kb fragment. (c1, d1) Western blot analyses of pH-modified virions showing the ∼17.5 kDa CP. (c2, d2) Coomassie blue-stained SDS-PAGE gels of pH-modified virions showing the ∼17.5 kDa CP and degradation products at pH 12. Numbers at the top of the figures are the treatment pH; ntc, negative template control; M, molecular size marker.

**Table 1.** pH effect on large segment amplifications using ToBRFV virions or virion RNA templates.

| Virions |  |  |  |  |  |  |  |  |  |  |  |  |
| --- | --- | --- | --- | --- | --- | --- | --- | --- | --- | --- | --- | --- |
| pH | 1.0 | 2.0 | 3.0 | 4.0 | 5.0 | 6.0 | 7.0 | 8.0 | 9.0 | 10.0 | 11.0 | 12.0 |
| *LS-PCR amplifications<br>/ total assays | 2/7 | 3/8 | 7/8 | 7/8 | 7/8 | 6/8 | 6/8 | 5/8 | 7/8 | 4/8 | 1/8 | 1/8 |
| Virion RNA |  |  |  |  |  |  |  |  |  |  |  |  |
| LS-PCR amplifications<br>/ total assays | 0/4 | 0/4 | 3/4 | 4/4 | 4/4 | 3/4 | 4/4 | 4/4 | 4/4 | 3/4 | 1/4 | 0/4 |

\*LS, large segment

### Extreme alkaline pH values promote degradation of ToBRFV coat protein

In addition to particle disassembly at pH 11-12 that was visualized by TEM (Fig.3 k, l), the pH-modified ToBRFV virions were subjected to a western blot analysis and Coomassie blue staining to determine the pH effects on ToBRFV CP. In the western blot, ToBRFV CP was detected by specific antibodies in all pH-modified ToBRFV virion preparations. However, at pH 12 very low CP levels were apparent. At pH 12, CP degradation products were observed in the Coomassie blue stained gel of the ToBRFV virions (Fig. 5d2).

### Soil treatment with alkaline solutions reduced ToBRFV soil-mediated infectious potential

Alkaline solutions in the pH range 8.35-13 were used for disinfection of ToBRFV-contaminated soil naturally occurring after a 4-6 month growth cycle of ToBRFV-infected tomato plants. The alkaline solutions were poured into the soil pots, saturating the soil for a 24 h incubation period, followed by soil remediation with tap water. Tomato seedlings with truncated roots planted in the remediated soil were tested for ToBRFV soil-mediated infection using ELISA. Soil treatment with either 3% Cl-TSP solution (pH 11.6) or with 3% Cl-TSP+1% KOH (pH 12.47) showed a significant reduction in infectious potential by soil-mediated ToBRFV, compared to untreated control (p<0.05; Two Sample t-Test assuming unequal variances) (Fig. 6a). Tomato plants grown in soil incubated with 3% Cl-TSP or 3% Cl-TSP+1% KOH showed the low infectivity of 6.3±6.3% and 5±5%, respectively, compared to the untreated control (infectivity of 39.7%±9.0). Tomato plants grown in soil incubated with tap water (pH 8.35) showed 19.1±17.7% infectivity. The high variability in the results of tap water effect hampers any prediction regarding the alkaline water efficiency as a soil disinfectant. Treatment with NaOH (pH 13) apparently caused ToBRFV degradation; however, soil structure was modified showing dispersion that led to plant mortality (Fig. 6a). The 3% Cl-TSP effect on soil disinfection was confirmed by testing ToBRFV CP content in the treated soil of one experiment using a western blot analysis (Fig. 6b). Tap water treatment in that experiment showed reduction in ToBRFV CP levels as well. The effect of Cl-TSP solutions on ToBRFV purified virions was also examined by TEM visualization and Coomassie blue staining following SDS-PAGE. Virion particles could not be visualized by TEM and virion CP could not be detected by the Coomassie blue-stained gel (Fig. 6d-f, g).

**Fig. 6.**
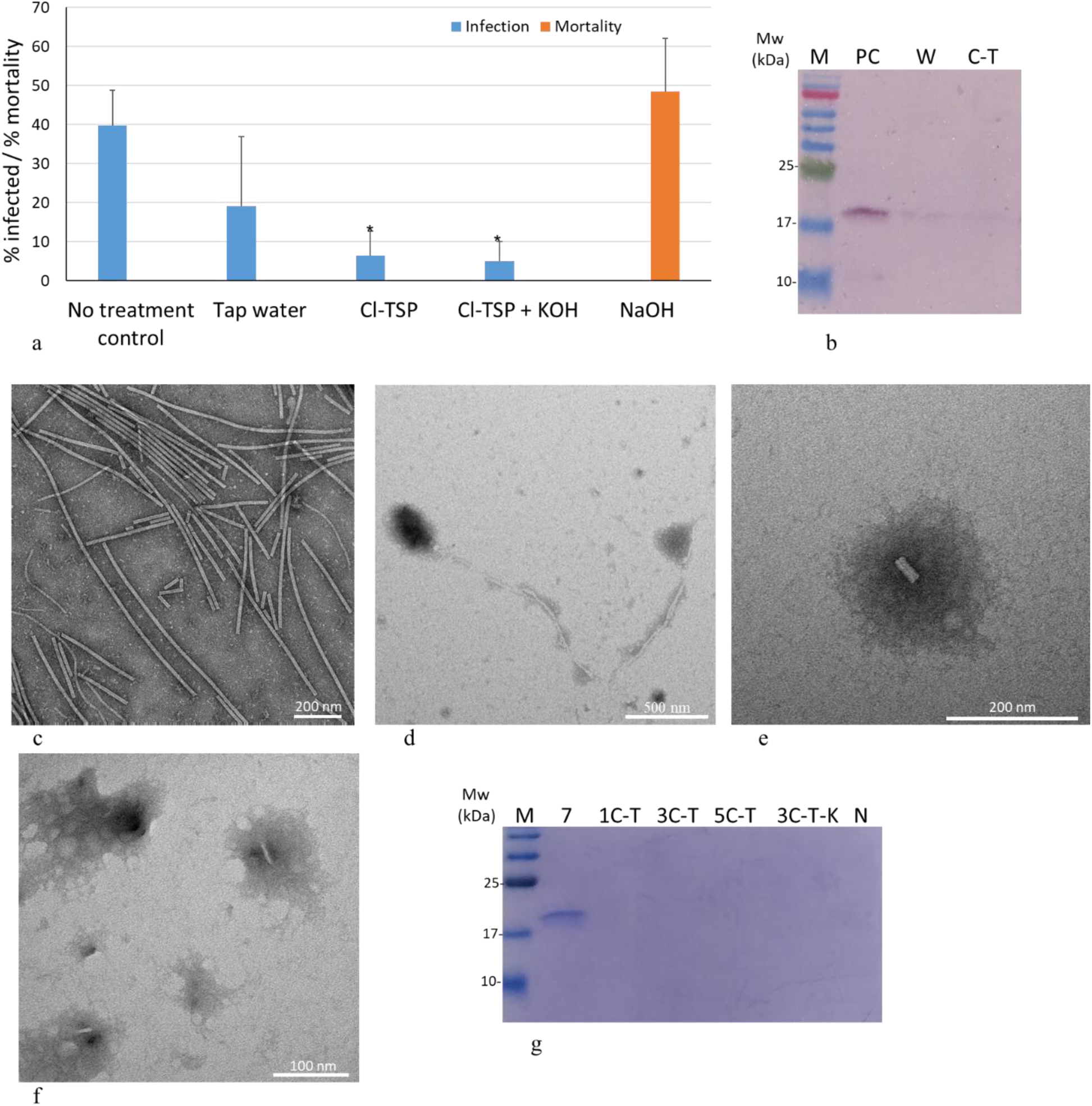
Soil disinfection with alkaline Cl-TSP solutions reduced ToBRFV soil-mediated infection of tomato seedlings and caused disintegration of ToBRFV virions. (a) ToBRFV soil-mediated infection of tomato seedlings following soil disinfection treatments with tap water Ph 8.35 (n=78), 3% Cl-TSP pH 11.6 (n=78), 3% Cl-TSP+1% KOH pH 12.47 (n=86) and NaOH pH 13 (n=81). Infectivity and mortality of tomato plants are presented as % ± SEM. Untreated soil served as a positive control (n=108). Asterisks represent a significant reduction p<0.05 (Two Sample t-Test assuming unequal variances) compared to the untreated positive control. (b) A western blot analysis of ToBRFV CP in 3% Cl-TSP treated soil. PC, positive control; W, tap water; C-T, 3% Cl-TSP. (c) A representative TEM micrograph of ToBRFV virions at pH 7 (d) A representative TEM micrograph of ToBRFV virions treated with 1% Cl-TCP. (e) A representative TEM micrograph of ToBRFV virions treated with 3% Cl-TCP; (f) A representative TEM micrograph of ToBRFV virions treated with 3% Cl-TCP+1% KOH. (g) A Coomassie blue stained gel of ToBRFV virions treated with soil disinfection solutions. 7, pH 7; 1C-T, 1% Cl-TSP pH 11.43; 3C-T, 3% Cl-TSP pH 11.6; 5C-T, 5% Cl-TSP pH 11.6; 3C-T-K, 3% Cl-TSP+1% KOH pH 12.47; N, NaOH pH 13; M, protein molecular size marker.

## Discussion

Soil-mediated tobamovirus infection of susceptible plants occurs by transmission through wounds in root tips, the virus reaching the upper parts of the plants (Allen, 1981; Broadbent, 1965; Pares et al., 1992). We have previously shown that the tobamovirus ToBRFV was transmitted *via* soil to susceptible tomato plants under two different study systems. The first was transmission from a viral inoculum poured into the planting pit an hour before planting. The second was transmission from naturally infected soil collected after a growth cycle of an infected crop (Dombrovsky et al., 2022; Klein et al., 2023). In both studies, root-truncated seedlings were planted to increase susceptibility to infection and increase test stringency. In the current study, we focused on testing ToBRFV longevity and soil-mediated infection under natural contamination conditions occurring after a 6-month growth cycle of contaminated crops, and tested infectivity using root-truncated tomato seedlings. In addition, we studied susceptibility of ToBRFV virions to treatments with acidic and alkaline solutions to promote soil remediation strategies that will reduce ToBRFV infectious potential in contaminated soil. For ToBRFV longevity studies, two earth piles kept either dry or wet were tested for ToBRFV infectious potential. The wet pile apparently increased decomposition of organic matter and we observed an increase in ToBRFV infectious potential until the 129^th^ day of pile assembly, whereas the dry earth pile showed reduction to 2% infectious potential on the 79^th^ day of pile-age. The decomposition of organic matter in the wet pile apparently allowed the elimination of ToBRFV infectivity by day 205 of pile-age (Fig. 2).

Our study of pH effects on ToBRFV particle stability showed that infectivity potential of ToBRFV was stable between the pH values of 2-10 tested on *N. glutinosa* plants (Fig. 4). These results differ from the results describing TMV susceptibility to pH treatments that showed particle stability over the pH range of 2-8 (Best & Samuel, 1936). There was also a report of a rapid inactivation of TMV below pH 3 and above pH 8 (Price, 1964; Stanley, 1946). However, a low pH 1 reduced infectivity potential of both TMV and ToBRFV tested on *N. glutinosa* plants (Fig. 4). At pH 1, TEM visualization of ToBRFV displayed reduced levels of rod-like particles and the amplification of a large genome segment by LS RT-PCR was not readily obtained (Fig 3a, Table 1). Importantly, at the high pH 9, TMV was partially inactivated, with the greatest fall in activity occurring in the first few minutes (Best & Samuel, 1936), implying that differences in incubation-time could not explain differences between TMV results and ToBRFV. At pH 9 and 10, TEM visualization of the pH-treated ToBRFV virions showed high concentrations of rod-like particles (Fig. 3i, j). In addition, the infectious potential of ToBRFV using *N. glutinosa* plants was not significantly different from that occurring at pH 7 and a large ToBRFV genome segment was readily amplified by LS RT-PCR (Figs. 4, 5a1, Table 1). These results also differ from studies of TMV CP disassembly that showed reduced bond strength at pH 9 (Caspar, 1964).

At the high pH 11, ToBRFV infectivity was lost, as tested on *N. glutinosa* plants, and TEM visualization showed truncated rod-like particles of up to ∼200 nm in length, along with discs resembling 20S discs formed by TMV CP at the lower pH 6.5-7 (Figs. 3k, 4) (Kegel & van der Schoot, 2006; Klug, 1999). As the pH is lowered, there was an increase in TMV CP rod length and at higher pH values, 4S aggregates were apparent (Bhyravbhatla et al., 1998). These data suggest that keeping the high pH values of tobamovirus solutions would prevent reformation of rod-like particles. In addition, studies of the TMV intracellular disassembly process in which carboxylate interactions were assumed to play a role, revealed that in alkaline solutions and low calcium ion concentrations, destabilization of the interaction between adjacent capsid subunits leads to viral disassembly (Caspar, 1964; Weis et al., 2019). At pH 12 no defined structures were visualized by TEM of ToBRFV virions and the ToBRFV CP was degraded (Figs. 3, 5d), which could prevent any potential reassembly.

For soil disinfection strategies, it is necessary to take into consideration virus attachment to various surfaces in the aquifer sediments, primarily controlled by electrostatic interactions (Loveland et al., 1996). The pH at the isoelectric point (pH_iep_) of the tobamovirus TMV was determined as ∼3.5 (Eriksson-Quensel & Svedberg, 1936). Accordingly, in solutions with pH values below 3.5 we could expect the tobamoviruses to be positively charged and most aquifer grain surfaces are positively charged below pH 4 (Loveland et al., 1996). However, we could not benefit from this repulsion between the tobamoviruses and most soil grain surfaces because at the acidic pH 1 there was a partial infectivity of ToBRFV observed on *N. glutinosa* plants and at pH 2 ToBRFV was highly infective (Fig. 4). Over the pH range 4-9 most aquifer grain surfaces are negatively charged (Loveland et al., 1996) and the tobamovirus would be negatively charged as well. However, the results of soil incubation with water at pH 8.35 showed low consistency of infectivity, suggesting that the electrostatic repulsion was not strong enough to promote ToBRFV displacement from soil.

To obtain soil treatment at pH ∼11 that compromised ToBRFV genome integrity and eliminated virion infectivity tested on *N. glutinosa* plants (Figs. 4, 5, Table 1) we used a 3% Cl-TSP solution (pH 11.6). In previous studies, soil treatment with 3% Cl-TSP reduced ToBRFV infectivity by inactivating a soil inoculum of ToBRFV (Dombrovsky et al., 2022; Klein et al., 2023). We now found that treatment of naturally contaminated soil with 3% Cl-TSP significantly reduced soil-mediated infection of tomato plants compared to the untreated positive control concomitant with reduction in ToBRFV CP levels (Fig. 6a, b). In a study of TSP activity on disruption of bacterial cell membranes the TSP effect was attributed to the solution pH values (Sampathkumar et al., 2003). However, our analyses of direct effect of Cl-TSP solutions at pH 11.6 on virion integrity visualized by TEM and Coomassie blue stained gel showed that virions and CP were undetectable, an effect more severe than the effect of the high pH 12 on ToBRFV virions (Figs. 3, 6). Therefore, a triple mechanism of disinfection is suggested to be attributed to Cl-TSP solutions: (i) the phosphate acts as a surfactant; (ii) the high pH causes protein degradation; (iii) the hypochlorite in aqueous solutions acts as an oxidizing and chlorinating agent.

To conclude: our data demonstrated that ToBRFV contaminated the soil following a growth cycle of naturally infected crops and the virus was highly stable and infectious at the pH range of natural waters (pH 4-9) (Loveland et al., 1996). To prevent soil-mediated ToBRFV infection, enhancement of soil disinfection could be provided by the use of alkaline disinfectant solutions or addition of alkaline solutions to other disinfection strategies such as steaming (Luvisi et al., 2015) or solarization.

## Acknowledgements

We would like to thank Dr. Victor Gaba for reviewing our manuscript. We acknowledge the contribution of Dr. Yael Friedman to TEM operation. We would like to thank Dr. Roman Sheinman for enlightening the issue of Cl-TSP chemical stability and Dr. Victoria Reingold for statistical analyses.

## Founding

SMART consortium, award by the Innovative Authority, Israel, grant number 500104773 and Horizon 2020, Virtigation consortium award number: 101000570.

## Author contributions

Conceptualization, A.D; Methodology O.M., E.S. N.L., E.B, O.L.; Validation, O.M, E.S., N.L. ; Formal Analysis, A.D. E.S., Investigation, A.D., M.R., O.M.; Resources, A.D., M.R.; Data Curation, O.M., A.D., E.S., ; Writing – Original Draft Preparation, O.M.; Writing – Review & Editing, E.S., A.D., M.R.; Visualization, O.M., E.S. ; Supervision, A.D., M.R. ; Project Administration, A.D.; Funding Acquisition, A.D.

## Competing interest

The authors declare no conflict of interest.

## Data availability

All data is presented in the manuscript.

## Notes

### Competing Interest Statement

The authors have declared no competing interest.

