## Supplementary material for "A study on tomato brown rugose fruit virus longevity in soil and virion susceptibility to various pH treatments contributes to optimization of a soil disinfection protocol": Fig.S1: Fig.S1-BioRxiv.pdf

**Fig. S1** Ionic strength of pH-modified ToBRFV virion and virion RNA solutions

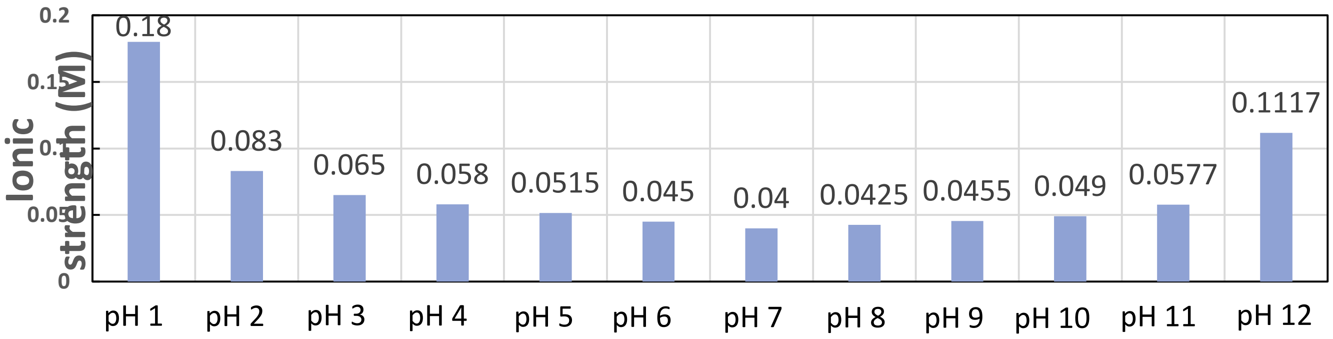
